# Differential effects of loss of *park7* activity on Iron Responsive Element (IRE) gene sets: Implications for the role of iron dyshomeostasis in the pathophysiology of Parkinson’s disease

**DOI:** 10.1101/2021.03.25.437102

**Authors:** Hui Yung Chin, Michael Lardelli, Lyndsey Collins-Praino, Karissa Barthelson

## Abstract

Mutation of the gene *PARK7* (*DJ1*) causes monogenic autosomal recessive Parkinson’s disease (PD) in humans. Subsequent alterations of PARK7 protein function lead to mitochondrial dysfunction, a major element in PD pathology. Homozygous mutants for the *PARK7*-orthologous genes in zebrafish, *park7*, show changes to gene expression in the oxidative phosphorylation pathway, supporting that disruption of energy production is a key feature of neurodegeneration in PD. Iron is critical for normal mitochondrial function, and we have previously used bioinformatic analysis of IRE-bearing transcripts in brain transcriptomes to find evidence supporting the existence of iron dyshomeostasis in Alzheimer’s disease. Here, we analysed IRE-bearing transcripts in the transcriptome data from homozygous *park7^−/−^* mutant zebrafish brains. We found that the set of genes with “high quality” IREs in their 5’ untranslated regions (UTRs, the HQ5’IRE gene set) was significantly altered in these 4-month-old *park7^−/−^* brains. However, sets of genes with IREs in their 3’ UTRs appeared unaffected. The effects on HQ5’IRE genes are possibly driven by iron dyshomeostasis and/or oxidative stress, but illuminate the existence of currently unknown mechanisms with differential overall effects on 5’ and 3’ IREs.

## Introduction

Parkinson’s disease (PD) is the second most common neurodegenerative disease, affecting approximately 1% of the population aged over 60 years. Most cases of PD are idiopathic, but a clear genetic link has been identified in approximately 5%-10% of PD cases (1). One gene, *PARK7*, implicated in autosomal recessive early-onset PD, encodes Parkinson disease protein 7 (PARK7). PARK7 protein has been suggested to act as a GSH-independent glyoxalase for detoxification of methylglyoxal and as a protein glycase responsible for restoring the function of proteins damaged by oxidative stress. However, these activities are disputed in PD (reviewed and analysed in (2)). PARK7 also has critical roles in maintaining mitochondrial function, sensing and responding to reactive oxygen species (ROS), and ultimately, acts in neuroprotection (reviewed in (3)).

PD is characterized by the specific depletion of substantia nigra pars compacta dopaminergic (SNc DA) neurons. These neurons make large numbers of synapses in the basal ganglia. Consequently, their high energy demands may make SNc DA neurons sensitive to energy deficiency (4). Many factors can affect energy production by the process of oxidative phosphorylation. In particular, ferrous iron (Fe^2+^) is incorporated into Fe-S clusters central to the function of the electron transport chain (ETC) in oxidative phosphorylation (5). ETC dysfunction causes oxidative stress that may lead to the (primarily) cytosolic PARK7 protein translocating into mitochondria to regulate the effects of reactive oxygen species (ROS) (5). This process is possibly altered in individuals with mutation of *PARK7*, leading to ETC dysfunction and dyshomeostasis of iron (via effects on Iron Regulatory Proteins, IRP1 & IRP2) and damaging dopaminergic neurons.

IRP1 and IRP2 bind IREs in the mRNAs of genes involved in iron homeostasis to regulate their translation and stability (reviewed in (6)). IRPs are regulated both by cellular ferrous iron status and oxidative stress (6). Previously, we defined sets of genes bearing IREs in either the 5’ or 3’UTRs of their transcripts (at lower or higher similarity to an IRE consensus sequence) in humans, mice and zebrafish (7). Using these, we found evidence supporting iron dyshomeostasis in Alzheimer’s disease (AD) brains, and in animal models of AD (7).

Orthologues of PD genes have previously been identified and manipulated in zebrafish. For example, Hughes et al. (8) developed a novel zebrafish model to examine the function of *PARK7.* RNA-seq was performed on whole brains from 4-month-old *park7^−/−^* zebrafish (compared to wild type), and gene set enrichment analysis (GSEA) was used to predict cellular processes altered by loss of *park7* function. Gene set enrichment profiles were observed consistent with disrupted metabolism and upregulation of genes associated with oxidative phosphorylation (8). We hypothesized that oxidative stress and/or iron dyshomeostasis in *park7^−/−^* zebrafish brains would alter binding of IRPs to transcripts containing IREs, thereby altering transcript stability. To explore this, we reanalysed the zebrafish brain transcriptome data from Hughes et al. to test for changes in the representation of IRE-containing gene sets in *park7^−/−^* zebrafish brains. We found that the HQ5’IRE gene set is significantly altered in 4-month-old *park7^−/−^* brains, while the HQ3’IRE gene set is not.

## Methods

To test for evidence of possible iron dyshomeostasis in *park7^−/−^* zebrafish brains, we performed enrichment analysis using *fry* (9) on the IRE gene sets (7). For detailed information of this re-analysis of Hughes et al. (8) data, see **Additional File 1**.

## Results

We previously defined sets of zebrafish genes according to whether these possess IRE-like motifs in either the 5’ or 3’ UTRs of their mRNAs, and whether their IREs match a canonical (high quality, HQ) or non-canonical IRE sequence (all) (7). We found that only transcripts of the HQ5’IRE gene set show statistically significant changes to gene expression as a group (**Figure 1A**) in 4-month-old *park7^−/−^* brains. Interestingly, the most upregulated gene of the HQ5’IRE gene set is *alas2.*

**Figure 1:**
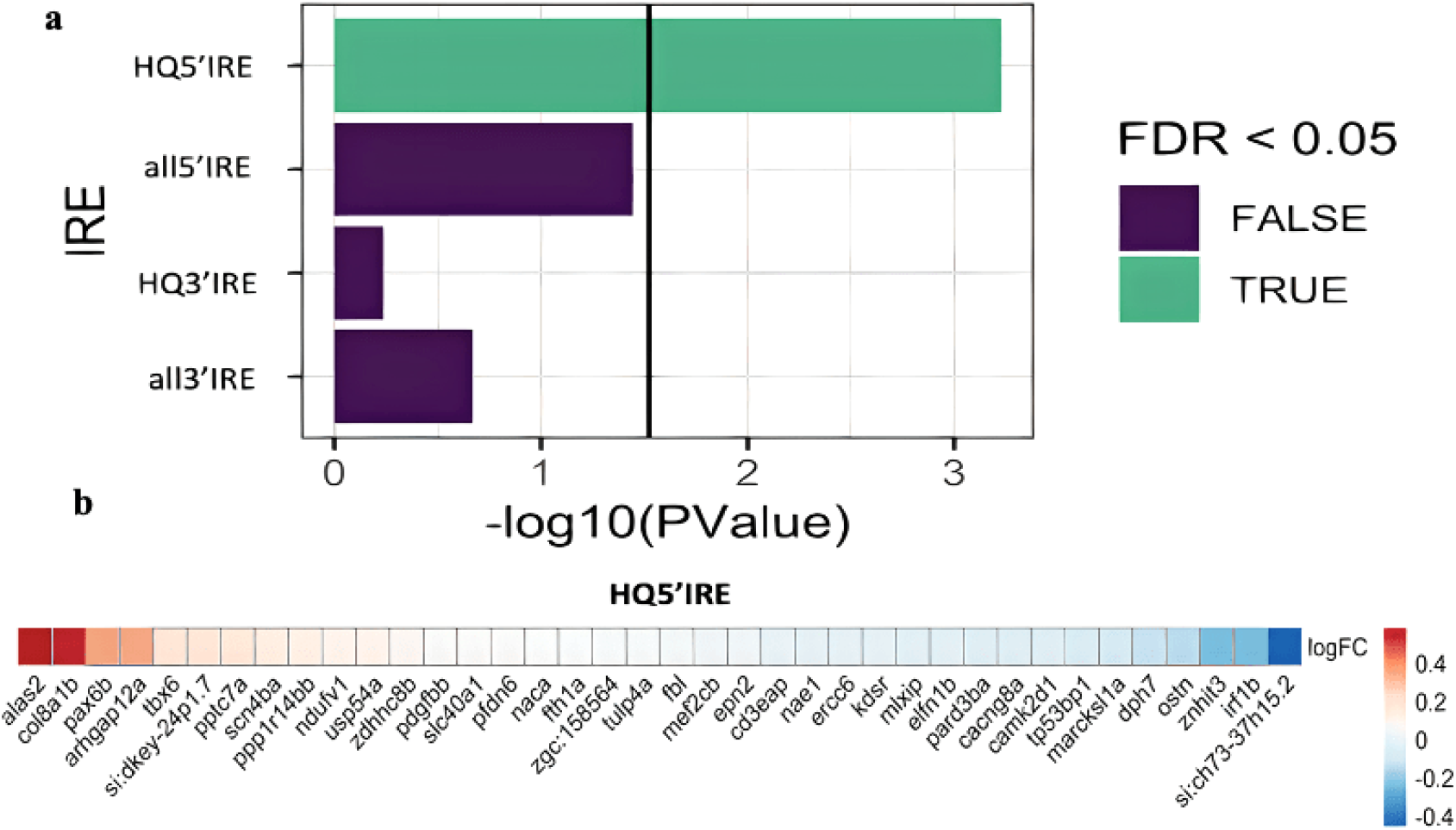
Enrichment analysis of IRE gene sets in *park7^−/−^* zebrafish brains. **a)** Bar plot showing the significance of enrichment of the four IRE gene sets in 4-month-old *park7^−/−^* zebrafish brains. The green bar shows that genes with high quality 5’IREs are significantly altered as a group. The vertical line indicates when the FDR-adjusted p-value = 0.05. **b)** Heatmap indicating the log2 fold change (logFC) of all HQ5’IRE genes detectable in the RNA-seq experiment.

## Discussion

Using our method to detect evidence for iron dyshomeostasis in RNA-seq data, we found highly significant alteration of the expression of genes with IREs in the 5’UTRs of their mRNAs in 4-month-old *park7^−/−^* brains.

Iron homeostasis is maintained by regulation of gene expression at several levels, including via transcription, mRNA stability, and mRNA translation (reviewed in (10)). The latter two phenomena are modulated by the iron regulatory proteins IRP1 and IRP2 when these bind to IREs. The highly significant alteration of expression of the HQ5’IRE gene set in 4-month-old *park7^−/−^* brains is likely due to changes in binding of IRP1 and/or IRP2 to IREs in the transcripts of these genes. However, since mutation of *PARK7* is known to cause oxidative stress (11) and oxidative stress can also affect IRP formation (reviewed in (6)), it is difficult to differentiate between oxidative stress and iron dyshomeostasis as contributing to changes in HQ5’IRE gene transcript abundance. Indeed, since iron is so important for mitochondrial function, iron dyshomeostasis and oxidative stress often co-occur (12). Interestingly, the HQ3’IRE gene set appeared unaffected in *park7^−/−^* brains and we currently have no explanation for why this should be so. However, it does point to the existence of mechanisms that can discriminate in binding of IRPs to IREs (or cause differences in the effects of such binding), depending on whether an IRE resides in the 5’ or 3’ UTR of a transcript.

Among members of the HQ5’IRE gene set, *alas2* transcript levels were observed to be increased in *park7^−/−^* brains. The relationship between *alas2* activity and IRPs is discussed in **Additional File 1**.

Taken together, our analysis of transcriptome data from *park7^−/−^* zebrafish brains supports the possibility of iron dyshomeostasis and/or oxidative stress as early preclinical events in PD. This provides impetus for further exploration using zebrafish of the effects of PD-linked genes on iron homeostasis and mitochondrial function, to provide mechanistic insight into PD for the development of therapeutics.

## List of abbreviations

PD: Parkinson’s disease
IRE: Iron responsive element
HQ: High quality
UTRs: untranslated
GSH: glutathione
SNc DA: substantia nigra pars compacta dopaminergic neurons
ETC: Electron transport chain
ROS: Reactive oxygen species
IRP1: Iron regulatory proteins 1
IRP2: Iron regulatory proteins 2
mRNA: Messenger ribonucleic acid
AD: Alzheimer’s disease
RNA-seq: RNA sequencing
GSEA: gene set enrichment analysis
*alas 2*: Delta-aminolevulinate synthase 2
FDR: false discovery rate

## Declarations

### Ethics approval and consent to participate

Not applicable

### Consent for publication

Not Applicable

### Availability of data and materials

The R code used to re-analyse the raw transcript counts of Hughes et al. can be found at https://github.com/karissa-b/dj1KO-RNAseq-IRE. The raw data from Hughes et al. is available from the Gene Expression Omnibus(GEO) database GSE135271 (https://www.ncbi.nlm.nih.gov/geo/query/acc.cgi?acc=GSE135271). The list of genes which contain an iron responsive element (IRE) in the untranslated regions of their mRNAs in zebrafish can be found at https://github.com/nhihin/ire.

### Competing interests

The authors declare that they have no competing interests.

### Funding

KB is supported by an Australian Government Research Training Program Scholarship and by funds from the Carthew Family Charity Trust. ML is an academic employee of the University of Adelaide. LCP is an academic employee of the University of Adelaide and is further supported by a Barbara Kidman Fellowship.

### Authors’ contributions

HYC drafted the manuscript. KB wrote the methodology, generated the diagrams and performed the bioinformatic analysis. ML, LCP and KB supervised and provided advice on the analysis of this paper. ML, KB, LCP and HYC edited the manuscript

## Acknowledgements

We would like to acknowledge the senior author of the paper (8), Mary Elizabeth Pownall for making available the raw transcripts data, and Nhi Hin for providing the sets of zebrafish genes containing iron-responsive elements.

## Notes

### Competing Interest Statement

The authors have declared no competing interest.

