## Additional File 1 for "Differential effects of loss of *park7* activity on Iron Responsive Element (IRE) gene sets: Implications for the role of iron dyshomeostasis in the pathophysiology of Parkinson’s disease"

### Supplemental Methods Description

We analysed the raw transcript counts of Hughes et al. using R (1). To obtain gene-level counts, we summed the number of counts arising from each transcript from a single gene. We then omitted genes which were undetectable. Genes were considered detectable if the log counts per million (logCPM) was > 1 in at least 3 out of the 6 samples. To test for iron dyshomeostasis, we obtained the lists of genes in zebrafish which contain IREs in the untranslated regions of their mRNAs (2). We restricted these gene sets to contain only the genes which were detected in the RNA-seq experiment. We then performed enrichment analysis using *fry* (3) with a mixed hypothesis. We considered an IRE gene set to be significantly altered if the false discovery rate (FDR)-adjusted p-value was < 0.05.

### Supplementary Discussion

Alas2 catalyses the first committed step of heme synthesis (4) that is important for the function of the ETC in mitochondria. Transcripts from *alas2* were the most upregulated among the detected genes of the HQ5’IRE gene set (log_2_ fold change of 0.59), although at a single-gene FDR- adjusted p-value far from statistical significance (0.75). In any case, caution is required in interpreting the meaning of this possible upregulation, as binding of IRPs to IREs in the 5’UTRs of mRNAs frequently inhibits their translation (as well as altering mRNA stability). Therefore, increased levels of *alas2* transcripts may, in this case, indicate decreased Alas2 protein synthesis, leading to decreased heme synthesis and disturbance of the ETC (which may cause additional oxidative stress).

Also, heme binds to heme regulatory motifs (HRMs) on both IRP1 and IRP2 to inhibit, by multiple mechanisms, their binding of IREs (5). If loss of *park7* function leads to increased oxidative stress, and that causes increased formation of IRP1 which blocks Alas2 protein synthesis, then the reduced production of heme might further increase IRP activity in a positive feedback loop. This would be expected to disturb iron homeostasis and be identifiable as changes in the stability of IRE-containing gene transcripts (as observed, at least, for the HQ5’IRE gene set).

Currently, we have no well supported explanation for why the HQ5’IRE gene set transcripts, as a group, should show significant changes in abundance while those of the 3’IRE gene sets do not, as this might require widespread differential binding of IRPs to IREs in 5’UTRs compared to IREs in 3’UTRs. However, it is notable that Zhu et al. (6) recently demonstrated that the ribosome recycling factor ABCE1 is sensitive to cellular iron deficiency and oxidative stress as it requires an Fe-S cluster for activity. Under iron deficiency and/or oxidative stress conditions, loss of ABCE1 activity causes failure of ribosomes to disassociate from mRNAs after stop codons so that they then interfere with the binding of proteins to motifs in 3’UTRs. Conceivably, this might also inhibit binding of IRPs to some 3’IREs, although likely only under extreme conditions, since 3’IREs have evolved to regulate transcripts in response to these conditions.
